# Three-dimensional Two-Photon Optogenetics and Imaging of Neural Circuits *in vivo*

**DOI:** 10.1101/132506

**Authors:** Weijian Yang, Luis Carrillo-Reid, Yuki Bando, Darcy S. Peterka, Rafael Yuste

## Abstract

We demonstrate a holographic system for simultaneous three-dimensional (3D) two-photon stimulation and imaging of neural activity in the mouse neocortex *in vivo* with cellular resolution. Dual two-photon excitation paths are implemented with independent 3D targeting for calcium imaging and precision optogenetics. We validate the usefulness of the microscope by photoactivating local pools of interneurons in awake mice visual cortex in 3D, which suppress the nearby pyramidal neurons’ response to visual stimuli.

---

To decipher how neural circuits function, it is important to precisely control the activity of specific neurons while simultaneously recording the activity of neuronal ensembles. Two-photon microscopy^1^ has proved its utility for *in vivo* calcium imaging because of its high selectivity, good signal-to-noise ratio, and depth penetration into scattering tissues^2^. It can be combined with two-photon optochemistry^3^ and optogenetics^4-8^ to allow for simultaneous readout and manipulation of neural activity with cellular resolution. But thus far, the combinations of both optical methods into an all-optical approach have been largely restricted to two-dimensional (2D) planes^3,5,6,8^ (but see Ref. ^9^). Since neural circuits are three dimensional, and genetically and functionally identified neuronal sub-populations are distributed throughout their volume, extending this method to three dimension (3D) appears essential to enable systematic studies of microcircuit computation and processing. Here we employed wavefront shaping strategy with a dual-beam two-photon microscope and simultaneously performed volumetric calcium imaging and 3D patterned photostimulations on mice cortex *in vivo*. We used a low repetition rate pulse-amplified laser for patterned stimulation, which significantly reduces the average laser power required for *in vivo* photoactivation, and minimizes thermal effects (supplementary notes).

We built a 3D microscope with two independent two-photon excitation lasers for imaging and photostimulation respectively (Fig. 1a). Each laser beam has control of its focus depth in the sample independent of objective movement, giving maximum flexibility and reducing possible mechanical perturbations to animal behavior. On the imaging path, we coupled a wavelength-tunable Ti:Sapphire laser to an electrically tunable lens (ETL)^10^ followed by a resonant scanner for high speed volumetric imaging. The ETL provides an adjustable axial focus shift up to 90 μm below and 200 μm above the objective’s nominal focal plane. On the photostimulation path, we used a spatial light modulator (SLM) to shape the wavefront, allowing flexible 3D beam splitting that targets the user defined positions in the sample, with an axial and lateral targeting error of 0.59±0. 54 μm and 0.82±0. 65 μm respectively across a 3D field of view (FOV) of 240x240x300 μm^3^ (Fig. 1b, 1c, supplementary Fig. S1; Methods). This SLM path was coupled through a pair of galvanometers to allow for fast sequential extension of the targeting FOV beyond the nominal addressable SLM-only range^11^. In optogenetics experiment, we also actuated this pair of galvanometric mirrors to spirally scan the focus over the cell body of each targeted neuron (see supplementary Fig. S2 for an exemplary 3D pattern with 100 targets on an autofluorescent plastic slide). Compared with alternative scanless strategy such as temporal focusing^6,9,12^, where the laser power is distributed across the whole cell body of each targeted neuron, this hybrid approach is simple, allowing a much smaller power budget, and scalable towards the amount of simultaneously targeted cells (supplementary notes). Though in this set of experiments, we imaged green fluorescence and photostimulated a red-shifted opsin, it is designed with switchable kinematic mirrors so that the lasers can be easily redirected to whichever path, and thus be utilized for red fluorophores and blue opsins.

**Figure 1.**
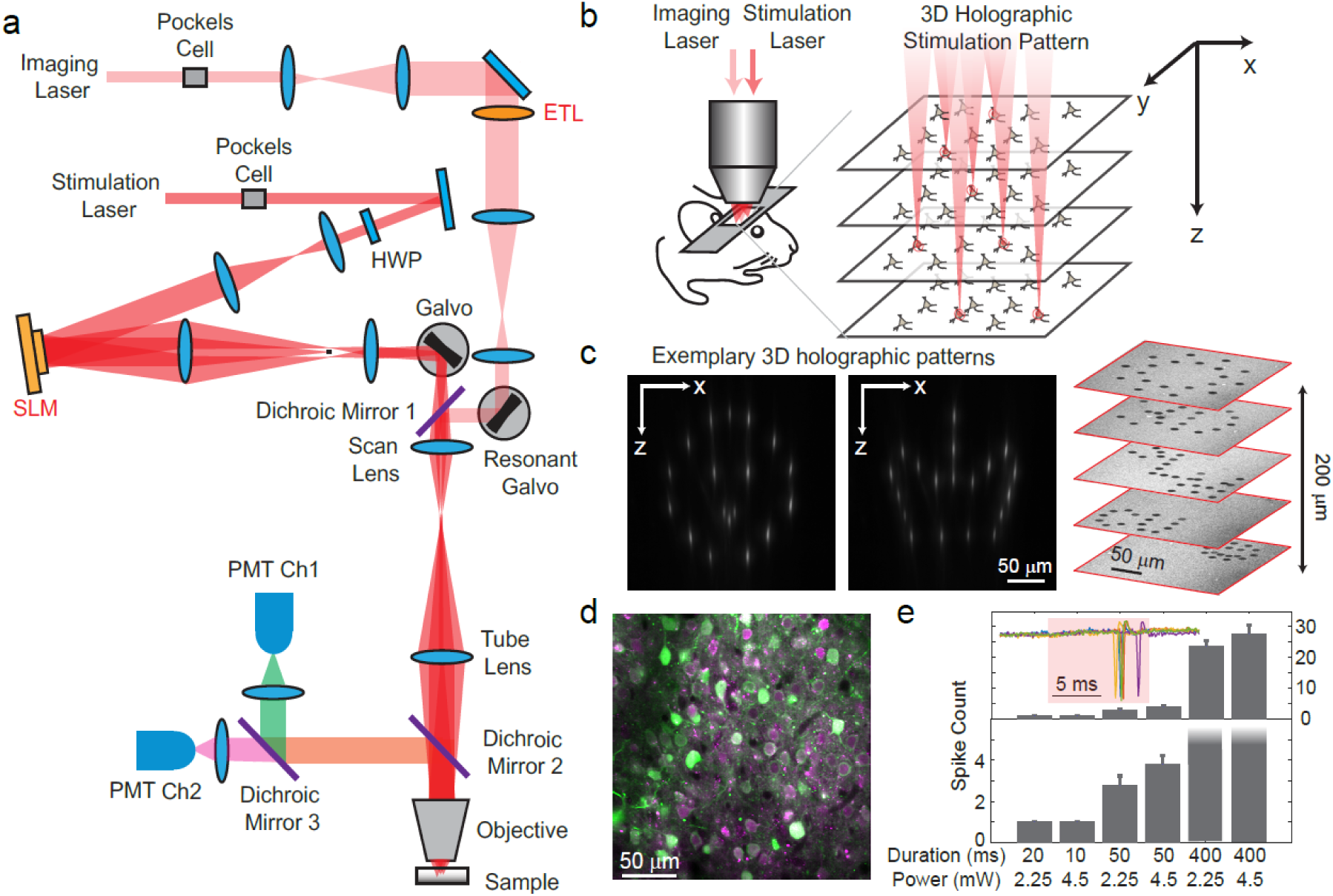
3D two-photon imaging and photostimulation microscope. (a) Dual two-photon excitation microscope setup. HWP, half-wave plate; SLM, spatial light modulator; ETL, electrically tunable lens; PMT, photomultiplier tube. (b) Schematics for simultaneous volumetric calcium imaging and 3D holographic patterned photostimulation in mouse cortex. (c) Exemplary 3D holographic patterns projected into Alexa 568 fluorescence liquid with its xz cross section captured by a camera (left and middle panel); 100 spots holographic pattern spirally scanned by a post-SLM galvanometric mirror bleaching an autofluorescence plastic slide (right panel). (d) A typical field of view showing neurons co-expressing GCaMP6s (green) and C1V1-mCherry (magenta). (e) Average spike counts (5 trials) of a neuron evoked by photostimulation with different spiral duration and average laser power. The inset shows the cell-attached recording of a 10 ms spiral stimulation over 5 trials. The red bar area indicates the photostimulation period.

We co-expressed the genetically encoded calcium indicator GCaMP6s^13^ and the red-shifted opsin C1V1^14^ into mouse primary visual cortex (V1, Fig. 1d), and excited them with 940 nm and 1040 nm light respectively. The separation of their excitation spectrum allowed for little cross-talk between the imaging and photostimulation beams (supplementary notes). C1V1-expressed cells were identified through a co-expressed mCherry fluorophore. Single spikes can be evoked with very low average laser power (∼2.25 mW with 20 ms spiral, or ∼4.5 mW with 10 ms spiral, 1 MHz pulse train, layer 2/3 *in vivo*, Fig. 1e), latency and jitter (12.2/7.4 ms latency, and 4.0/2.3 ms jitter for the two conditions, supplementary Fig. S3).

We tested our 3D all-optical system by targeting and photoactivating selected pyramidal cells across layer 2/3 of mouse V1 *in vivo* (mouse in anesthetized condition), while simultaneously monitoring the neuronal activity in three selected planes (240x240 μm^2^ FOV for each plane) at 6.67 vol/s. Neurons were photoactivated one at a time (supplementary Fig. S4), or as groups (*M* neurons simultaneously, *M*=3∼27, Fig. 2) and the majority of the targeted cells (86%±6%, Methods) showed clear calcium transients in response to the photostimulation (Fig. 2b-c).

**Figure 2.**
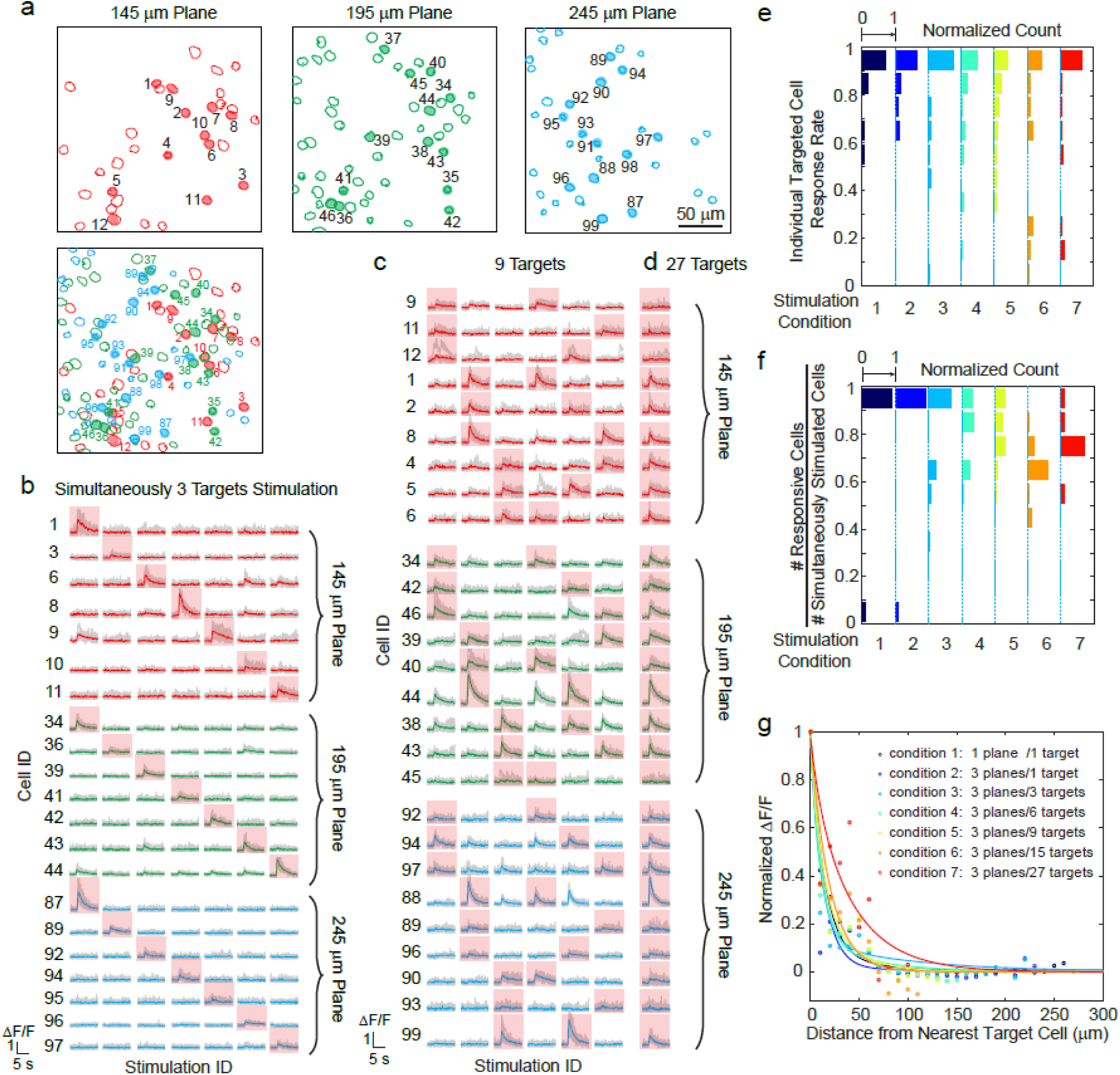
Simultaneous photostimulation of pyramidal cells in layer 2/3 of mouse V1 *in vivo*. (a) Contour maps showing the spatial location of the cells in three individual planes (top; 145 μm, 195 μm, and 245 μm from pial surface), and their 2D overlap projection (bottom). Cells with shaped color are the targeted cells. (b)-(d) Representative photostimulation triggered calcium response of the targeted cells and non-targeted cells, for different stimulation patterns. A total number of (b) 3, (c) 9, and (d) 27 cells across three planes were simultaneously photostimulated. (e) Histogram of individual targeted cell response rate in different stimulation conditions. The stimulation conditions are listed in (g). (f) Histogram of the percentage of responsive cells in a targeted group in different stimulation conditions. (g) Response of the non-targeted cells to the photostimulation versus distance to their nearest targeted cell. ΔF/F is normalized to the averaged response of the targeted cells.

We investigated the reliability of the photoactivation and its effect on non-targeted cells. We performed 8∼11 trials for each stimulation pattern. Cells not responding to photostimulation under any condition were excluded in this analysis (see Methods). As *M* increased, the response rate of individual targeted cell remained high and stable (Fig. 2e, 82%±9%). Within a group of cells that were simultaneously photostimulated, the percentage of responsive cells was also stable (Fig. 2f, 82%±9%). Although we had high targeting accuracy and reliability for exciting targeted cells, we also observed occasional activity in non-targeted cells (unspecific activation) during photostimulation. This was distance-dependent, and as the distance *d* between the non-targeted cells and their nearest targeted cells decreased, their probability of activation increased (Fig. 2g). And, for the same *d*, this probability increased with *M*. Nevertheless, it was confined (half response rate) within *d*<25 μm in all conditions (*M*=3∼27 across 3 planes spanning a volume of 240x240x100 μm^3^), in spite of the fact that we specifically used extremely long spiral durations (∼480 ms) to maximally emulate an undesirable photostimulation scenario. The activation of the non-targeted cells could occur through different mechanisms, such as by direct stimulation (depolarization) of the cells through their neurites that course through the photostimulation region, or by an indirect one, through synaptic activations by targeted cells, or a combination of the two. Direct stimulation could be reduced with somatic-restricted expression^15^, as well as sparse expression.

For 3D activation, one particularly attractive target are interneurons. Different interneuron classes participate in cortical microcircuits that could serve as gateways for information processing^16,17^. These interneurons are located sparsely in the cortex, and are highly connected to excitatory populations, and are known to strongly modulate activity. However, the relative effects of simultaneous stimulation of a selective subset of interneurons with single cell resolution has never been reported, as previous studies have largely relied on one-photon optogenetics where widespread activation is the norm^18,19^ (but see Ref. ^20^ for single cell stimulation). We used our all-optical approach to examine the effect of photoactivating a specific sub-pool of interneurons in 3D on the activity of pyramidal cells that responded to visual stimuli in an awake head-fixed mouse (Fig. 3).

**Figure 3.**
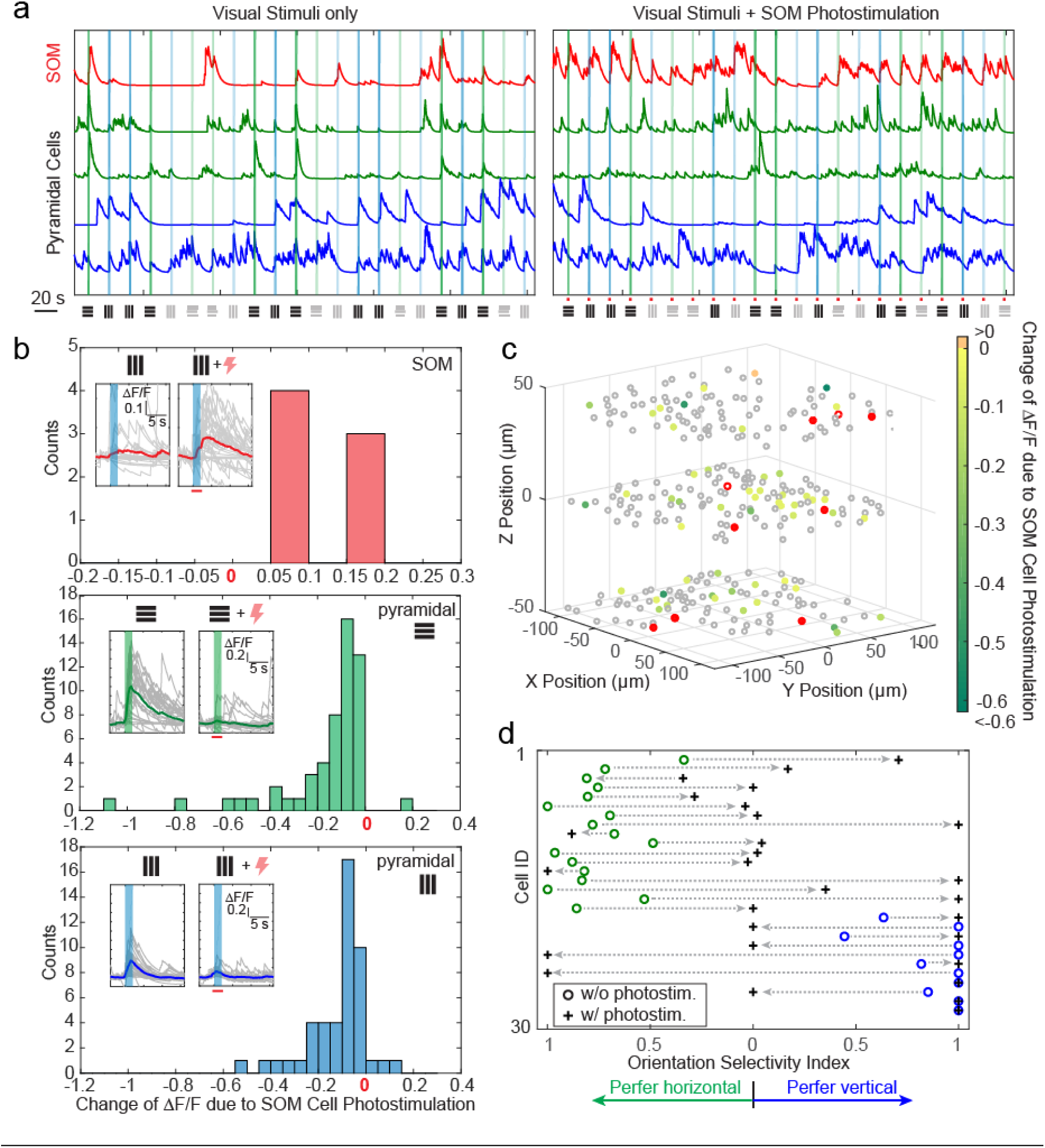
Simultaneous photostimulation of SOM interneurons suppresses responses to visual stimuli of pyramidal cells in awake mice. (a) Normalized calcium traces (ΔF/F) of representative targeted SOM cells and pyramidal cells that are responding to visual stimuli, without (left panel) and with (right panel) SOM cell photostimulation. The normalization factor for each cell stays the same across the two conditions. The shaped regions indicate the visual stimuli period. (b) Histogram of ΔF/F change for different cell populations that show significant activity change (p<0.05) due to SOM cell photostimulation (*M*=9). Top panel, targeted SOM cells (7 out of 9 show significant responses to photostimulation). Middle and bottom panels, pyramidal cells responding to horizontal or vertical drifting-gratings respectively. The inset compares the activity of a representative cell without and with targeted SOM cell photostimulation; the shaped regions indicate the visual stimuli period. (c) Spatial map of all the recorded cells. Pyramidal cells responding to horizontal drifting-gratings and showing significant ΔF/F change due to SOM cell photostimulation [cell population in the middle panel of (b)] are color coded according to their ΔF/F change. The targeted SOM cells are outlined in red, and those responding are shaped in red. (d) Orientation selectivity index of a group of pyramidal cells without and with SOM cell photostimulation. This group of cells were selected if they show significant response and orientation preference to visual stimuli without SOM cell photostimulation, and show significant change of ΔF/F during SOM cell photostimulation (p<0.05).

We drove Cre-dependent C1V1 in somatostatin (SOM) inhibitory interneurons (SOM-Cre mice), while simultaneously expressing GCaMP6s in both pyramidal cells and interneurons, in layer 2/3 of mouse V1. We first imaged the responses of pyramidal cells across 3 planes (separated by ∼45 μm each) to orthogonal visual stimuli consisting of drifting grating without photostimulation. We then simultaneously photostimulated a group of SOM cells (*M*=9, with 7 showing significant responses) across these 3 planes concurrently with the visual stimuli (Methods). We observed a significant suppression in response among 46% and 35% of the pyramidal cells that originally responded strongly to the horizontal and vertical drifting-grating respectively. The orientation selectivity index (OSI) for highly selective cells was also altered by the SOM cell photoactivation. This is consistent with previous reports showing that SOM cells inhibit nearby pyramidal cells with one-photon optogenetics *in vivo*^18,19^ or with two-photon uncaging *in vitro*^21^. On the other hand, our two-photon approach provides high precision 3D manipulation over groups of cells, and simultaneous readout of neuronal activity across the network *in vivo*. Thus, our approach could be useful to dissect the excitatory and inhibitory interactions in cortical circuits *in vivo*.

Our 3D all-optical method could be used to study cell connectivity, ensemble organization and information processing in neural circuits. It extends previous methods from a 2D plane to a 3D volume, representing an important step of precision optogenetics towards large spatial scales. The high-peak power of the low-repetition rate photostimulation laser allows targeting a large group of cells with low average power (e.g. 27 target cells in layer 2/3 with 86 mW in total). We choose an ETL for volumetric imaging because of its low cost and great compatibility with most existing two-photon microscopes. One of the future directions could be replacing the ETL with another SLM to perform multiplane imaging^22^ and adaptive optics, which could increase the frame rate and improve the imaging quality. While we use a relatively low excitation NA (∼0.35) beam that is limited by the small mirror size (3 mm) of the post-SLM galvanometric scanners, increasing the mirror size is a straightforward future development of this method that would increase this NA, and decrease the axial point spread function (currently 14.5 μm full-width-at-half-maximum, supplementary Fig. 1a). It would also improve the axial resolution of the spiral photostimulation (currently ∼20 μm, measured by displacing the 12 μm diameter spiral pattern relative to the targeted neuron, supplementary Fig. S3), and thus reduce the unspecific activation of the non-targeted cells. Another approach to suppress the unspecific activation is to use a somatic-restricted opsin. A somatic-restricted channelrhodopsin 2 was reported recently^15^, and showed reduced activation on non-targeted cells *in vitro*. While we demonstrated the successful manipulation of the targeted neural microcircuits in awake head-fixed behaving mice by photostimulating a targeted group of interneurons, we expect this 3D all-optical method would find its many other applications in dissecting the neural circuits.

## METHODS

### Microscope design

The optical setup is illustrated in Fig. 1a, which is composed of two femtosecond pulse lasers and a custom-modified two-photon laser scanning microscope (Ultima In Vivo, Bruker Corporation). The laser source for imaging is a pulsed Ti:sapphire laser (Coherent, Chameleon Ultra II). Its wavelength is tuned to 940 nm for GCaMP6s imaging or 750 nm for mCherry imaging respectively. The laser power is controlled with a Pockels cell (Conoptics, 350-160-BK, 302RM controller). The laser beam is expanded by a 1:3.2 telescopes (f=125 mm and f=400 mm) and coupled to an electrically tunable lens (Optotune Switzerland AG, EL-10-30-C-NIR-LD-MV) with a clear aperture of 10 mm in diameter. The transmitted beam is rescaled by a 3.2:1 telescope (f=400 mm and f=125 mm) and imaged onto a resonant scanner and galvanometric mirror, both located at the conjugate planes to the microscope’s objective pupil. The beam is further scaled by a 1:1.33 telescope before coupled into a scan lens (f=75 mm), a tube lens (f=180 mm) and the objective lens (Olympus 25x N.A. 1.05 XLPlan N), yielding an excitation NA ∼ 0.45. The fluorescence signal from the sample is collected through the objective lens and split at a dichroic mirror (Chroma Technology, HQ575dcxr, 575 nm long pass) to be detected in two bi-alkali photomultiplier tubes, one for each wavelength range. Two different bandpass filters (Chroma Technology, 510/20-2P, and 607/45-2P) are placed in front of the corresponding PMT respectively.

The optical path for the photostimulation is largely independent from the imaging, except that they combine at a dichroic mirror (Chroma Technology, T1030SP, 1030 nm short pass) before the scan lens, and thus share the subsequent optical path. The laser source for photostimulation is a low repetition rate (200 kHz∼1 MHz) pulse-amplified laser (Spectra-physics, Spirit 1040-8), operating at 1040 nm wavelength. Its power is controlled by a Pockels cell (FastPulse Technology, 1147-4-1064). A λ/2 waveplate (Thorlabs, AHWP05M-980) is used to rotate the laser polarization so that it is parallel to the active axis of the spatial light modulator (Meadowlark Optics, HSP512-1064; 7.68x7.68 mm^2^ active area, 512×512 pixels). The beam is expanded by two telescopes (1:1.75, f=100 mm and f=175 mm; 1:4, f=50 mm and f=200 mm) to fill the active area of the SLM. The reflected beam from the SLM is scaled by a 3:1 telescope (f=300 mm and f=100 mm) and imaged onto a set of close-coupled galvanometer mirrors, located at the conjugate plane to the microscope’s objective pupil. A beam block made of a small metallic mask on a thin pellicle is placed at the intermediate plane of this telescope to remove the zero-order beam. The photostimulation laser beam reflected from the galvanometer mirrors are then combined with the imaging laser beam at the 1030 nm short pass dichroic mirror.

The imaging and photostimulation is controlled by a combination of PrairieView (Bruker Corporation) and custom software running under MATLAB (The Mathworks) on a separate computer. The Matlab program was developed to control the ETL through a data acquisition card (National Instrument, PCIe-6341) for volumetric imaging, and the SLM through PCIe interface (Meadowlark Optics) for holographic photostimulation. The two computers are synchronized with shared triggers. At the end of each imaging frame, a signal is received to trigger the change of the drive current (which is converted from a voltage signal from the data acquisition card by a voltage-current converter [Thorlabs, LEDD1B]) of the ETL, so the imaging depth is changed for the following frame. The range of the focal length change on sample is ∼ +90 μm ∼ −200 μm (“+” means longer focal length). The intrinsic imaging frame rate is ∼30 fps with 512 x 512 pixel image. The effective frame rate is lower as we typically wait 10∼17 ms in between frames to let the ETL fully settle down at the new focal length. The control voltage of the Pockels cell is switched between different imaging planes to maintain image brightness.

### SLM hologram and characterization

The phase hologram on the SLM, ø(*u*, *v*), can be expressed as:

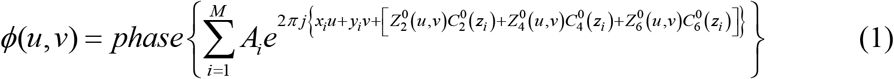

where [*x*_*i*_, *y*_*i*_, *z*_*i*_] (*i*=1,2…*M*) is the coordinate of the cell body centroid (*M* targeted cells in total), and *A*_*i*_ is the electrical field weighting coefficient for the *i*^th^ target (which controls the laser power it receives). 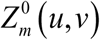 and 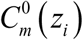 are the Zernike polynomials and Zernike coefficients, respectively, which sets the defocusing and compensates the first-order and second-order spherical aberration due to defocusing. Their expressions are shown in the table below. The hologram can also be generated by 3D Gerchberg-Saxton algorithm, but it takes extra steps to incorporate spherical aberration compensation. We adapt Eq. (1) as a simpler method.

|  |  |
| --- | --- |
| Defocus |  |
| Zernike polynomials | $Z_2^0(u, v) = \sqrt{3} [2(u^2 + v^2) - 1]$ |
| Zernike coefficients | $C_2^0(z) = \frac{nkz \sin^2 \alpha}{8\pi\sqrt{3}} \left( 1 + \frac{1}{4} \sin^2 \alpha + \frac{9}{80} \sin^4 \alpha + \frac{1}{16} \sin^6 \alpha + \dots \right)$ |
| First-order spherical aberration |  |
| Zernike polynomials | $Z_4^0(u, v) = \sqrt{5} [6(u^2 + v^2)^2 - 6(u^2 + v^2) + 1]$ |
| Zernike coefficients | $C_4^0(z) = \frac{nkz \sin^4 \alpha}{96\pi\sqrt{5}} \left( 1 + \frac{3}{4} \sin^2 \alpha + \frac{15}{18} \sin^4 \alpha + \dots \right)$ |
| Second-order spherical aberration |  |
| Zernike polynomials | $Z_6^0(u, v) = \sqrt{7} [20(u^2 + v^2)^3 - 30(u^2 + v^2)^2 + 12(u^2 + v^2) - 1]$ |
| Zernike coefficients | $C_6^0(z) = \frac{nkz \sin^6 \alpha}{640\pi\sqrt{7}} \left( 1 + \frac{5}{4} \sin^2 \alpha + \dots \right)$ |
| $n$, refractive index of media between the objective and sample; $k$, the wavenumber; $z$, the axial shift of the focus plane in the sample; $u, v$, coordinates on the SLM phase mask; $n \sin \alpha$, the NA of the objective. | |

To match the defocusing length set in SLM with the actual defocusing length, we adjusted the “effective N.A.” in the Zernike coefficients following the calibration procedure described in Ref. ^22^. To register the photostimulation beam’s targeting coordinate in lateral directions, we projected 2D holographic patterns to burn spots on the surface of an autofluorescent plastic slide and visualized them by the imaging laser. An affine transformation can be extracted to map the coordinates. We repeated this registration for every 25 μm defocusing depth on the sample, and applied a linear interpolation to the depths in between. To characterize the lateral registration error, we actuated the SLM and burned spots on the surface of an autofluorescent plastic slide across a field of view of 240 μm x 240 μm with a 7x7 grid pattern. We then imaged the spots pattern with the imaging laser and measured the registration error. This was repeated for different SLM focal depths. An alternative method to register the targeting coordinate is to set the photostimulation laser in imaging mode, actuate the SLM for different lateral deflection, and extract the transform matrix from the acquired images and that acquired from the imaging laser. To characterize the axial registration error, we used the photostimulation laser to image a slide with quantum dots sample. The SLM was set at different focal depths, and a z-stack was acquired for each setting to measure the actual defocus and thus the axial registration error. In all these registration and characterization procedures, we used water as the media between the objective and the sample, and we kept the focus of the photostimulation laser at the sample surface by translating the microscope stage axially. We note that the refractive index of the brain tissue is slightly different from that of water (∼2%), and this could cause an axial shift of the calibration. This could be corrected in the Zernike coefficients. In practice, we found this effect is negligible, as the typical focal shift by the SLM is relatively small (<150 μm) and the axial PSF is large.

Due to the chromatic dispersion and finite pixel size of SLM, the SLM’s beam steering efficiency drops with larger angle, leading to a lower beam power for targets further away from the center field of view (in xy), and nominal focus (in z). The characterization result is shown in supplementary Fig. S1. A linear compensation can be applied in the weighting coefficient *A*_*i*_ in Eq. (1) to counteract this non-uniformity. In practice, these weighting coefficients can be adjusted such that the targeted neurons show clear response towards photostimulation.

### Animals and surgery

All experimental procedures were carried out in accordance with animal protocols approved by Columbia University Institutional Animal Care and Use Committee. Multiple strain of mice were used in the experiment, including C57BL/6 wild-type and SOM-cre (The Jackson Laboratory, stock no. 013044) mice at the age of postnatal day (P) 45-120. Virus injection was performed to layer 2/3 of the left V1 of the mouse cortex, 3∼8 weeks prior to the craniotomy surgery. For the C57BL/6 wild-type mice, virus AAV1-syn-GCaMP6s and AAVDJ-CaMKII-C1V1-(E162T)-TS-p2A-mCherry-WPRE was mixed and injected for calcium imaging and photostimulation; virus AAV8-CaMKII-C1V1-p2A-EYFP was injected for electrophysiology. For the SOM-cre mice, virus AAV1-syn-GCaMP6s and AAVDJ-EF1a-DIO-C1V1-(E162T)-p2A-mCherry-WPRE was mixed and injected. The virus was front-loaded into the beveled glass pipette (or metal pipette) and injected at a rate of 80∼100 nl/min. The injection sites were at 2.5 mm lateral and 0.3 mm anterior from the lambda, putative monocular region at the left hemisphere.

After 3∼8 weeks of expression, mice were anesthetized with isoflurane (2% by volume, in air for induction and 1-1.5% during surgery). Before surgery, dexamethasone sodium phosphate (2 mg per kg of body weight; to prevent cerebral edema) were administered subcutaneously, and enrofloxacin (4.47 mg per kg) and carprofen (5 mg per kg) were administered intraperitoneally. A circular craniotomy (2 mm in diameter) was made above the injection cite using a dental drill. A 3-mm circular glass coverslip (Warner instruments) was placed and sealed using a cyanoacrylate adhesive. A titanium head plate with a 4 mm by 3.5 mm imaging well was attached to the skull using dental cement. After surgery, animals received carprofen injections for 2 days as post-operative pain medication. The imaging and photostimulation experiments were performed 1∼14 days after the chronic window implantation. During imaging, the mouse is either anesthetized with isoflurane (1-1.5% by volume in air) with a 37°C warming plate underneath or awake and can move freely on a circular treadmill with its head fixed.

### Visual stimulation

Visual stimuli were generated using MATLAB and the Psychophysics Toolbox^23^ and displayed on a monitor (Dell; P1914Sf, 19-inch, 60-Hz refresh rate) positioned 15 cm from the right eye, at 45° to the long axis of the animal. Each visual stimulus session consisted of four different trials, each trial with a 2 s drifting square grating (0.04 cycles per degree, two cycles per second), followed by 18 s of mean luminescence gray screen. Four conditions (combination of 10% or 100% grating contrast, 0° or 90° drifting grating direction) were presented in random order in the four trials in each session.

### Photostimulation parameters

The photostimulation laser beam is split into multiple foci, and spirally scanned (∼12 μm final spiral diameter, 8∼32 rotations) by a pair of post-SLM galvanometric mirror over the cell body of each targeted cell. For neurons in layer 2/3 of mice V1, the typical average power used for each spot is 2 mW∼ 5 mW (1 MHz pulse repetition rate). The spiral duration ranged from 10 ms to 1000 ms. We specifically used extremely long spiral durations (∼480 ms) when studying the photostimulation effect on the non-targeted cells (Fig. 2) to emulate an undesirable photostimulation scenario. In the experiment that the SOM cells were photostimulated when the mouse were receiving visual stimuli, the photostimulation started 0.5 s before the visual stimuli, and ended 0.3 s after the visual stimuli finished. Since the visual stimuli lasted for 2 sec, the photostimulation lasted for 2.8 sec. This long photostimulation was composed of 175 continuous spiral scans, each lasting 16 ms.

### Data analysis

The recording from each plane was first extract from the raw imaging files, followed by motion correction using a pyramid approach^24^ or fast Fourier transform-based algorithm^25^. A constrained nonnegative matrix factorization (CNMF) algorithm^26^ was used to extract the fluorescence traces (ΔF/F) of the region of interested (i.e. neuron cell bodies in the field of view). The CNMF algorithm also outputs a temporally deconvolved signal, which is related to the firing event probability. The ΔF/F induced by the photostimulation was quantified with the mean fluorescence change during the photostimulation period over the mean fluorescence baseline within a 2 sec window before the photostimulation.

To detect the activity events from each recorded neuron, we thresholded the temporally deconvolved ΔF/F signal with at least 2 standard derivations from the mean signal. Independently, a temporal first derivative is applied to the ΔF/F trace. The derivative is then threshold at least 2 standard derivations from the mean. At each time point, if both are larger than the threshold, an activity event is recorded in binary format.

A cell is determined as not responding to photostimulation if there is no single activity event detected or no typical action-potential-corresponding calcium transient during photostimulation period for multiple trials. These non-responding cells could be due to a poor expression of C1V1.

The orientation selectivity index and preference of the visual stimuli is calculated as the amplitude and sign of (ΔF/F|_90_ − ΔF/F|_0_) / (ΔF/F|_90_ + ΔF/F|_0_) respectively, where ΔF/F|_90_ and ΔF/F|_0_ is the mean ΔF/F during the visual stimuli with 90o and 0o drifting grating respectively.

In general, the GCaMP6s should not be over expressed; which could otherwise cause a fluorescence background during photostimulation (supplementary notes). This background could superimpose onto the calcium imaging data. Since the pixel rate (∼8.2 MHz) of the calcium imaging is much faster than the photostimulation laser’s pulse repetition rate (1 MHz), this background artifact appears to be a mesh grid shape in the calcium imaging movie (supplementary Fig. S5). Typically it is small and does not impact the above data analysis. In the case that it is strong, the recorded frames during photostimulation is pre-processed to suppress this background artifact (supplementary Fig. S5). To detect the pixels having this artifact, we consider both their fluorescence value and their geometry. First we detect candidate pixels by identifying pixels whose value is significantly higher from the average value calculated from a few frames just before and just after the stimulation. Second, these candidate pixels are tested for connectedness within every horizontal and vertical line of each frame, and the width of the connections compared to that expected based on the stimulation condition. If both these conditions hold, these pixels are marked as “contaminated” and the fluorescence value at these pixels during the stimulation are replaced by those in their adjacent “clean” pixels. This procedure was used for the data in Fig. 3. This pre-processing significantly suppresses the artifacts while maintaining the original signal. Nevertheless, to avoid any analysis bias, we further approximated the neuronal response by using the ΔF/F signal just after the photostimulation, when there was no background artifact. The same analysis procedure was implemented to the control experiment when there was no photostimulation.

### In vivo electrophysiological recordings

Mice were head-fixed and anaesthetized with isoflurane (1.5∼2%) throughout the experiment. Dura was carefully removed in the access point of the recording pipette. 2% agarose gel in HEPES-based artificial cerebrospinal fluid (ACSF) (150 mM NaCl, 2.5 mM KCl, 10 mM HEPES, 2 mM CaCl_2_, 1 mM MgCl_2_, pH was 7.3) was added on top of the brain to avoid movement artifacts. Patch pipettes of 5∼7 MΩ pulled with DMZ-Universal puller (Zeitz-Instrumente Vertriebs GmbH) were filled with ACSF containing 25 μM Alexa594 to visualize the tip of the pipettes. C1V1-expressing cells were targeted using two-photon microscopy *in vivo*. During recordings, the space between the objective and the brain was filled with ACSF. Cell-attached recordings were performed using Multiclamp 700B amplifier (Molecular Devices), in voltage-clamp mode. Sampling rate was 10 kHz, and the data was low-pass filtered at 4 kHz using Bessel filter.

## ACKNOWLEDGMENTS

This work is supported by the NEI (DP1EY024503, R01EY011787), NIMH (R01MH101218, R41MH100895, R01MH100561, R44MH109187), and DARPA contracts W91NF-14-1-0269 and SIMPLEX N66001-15-C-4032. This material is based upon work supported by, or in part by, the U.S. Army Research Laboratory and the U.S. Army Research Office under contract number W911NF-12-1-0594 (MURI). W. Yang holds a career award at the scientific interface from Burroughs Wellcome Fund. Y. Bando holds a fellowship from Uehara Memorial Foundation. The authors thank Reka Letso, Mari Bando, and Azi Hamzehei for virus injection of the mice; Sean Quirin for the initial software for SLM control; Adam Packer and Alan Mardinly for fruitful discussions.

### AUTHOR CONTRIBUTIONS

W.Y., L.C.-R., D.S.P. and R.Y. designed the study; W.Y. and D.S.P. designed and built the hardware and software of the holographic stimulation and volumetric imaging. W.Y. and Y. B. performed the experiments. W.Y. performed the analysis. W. Y. wrote the original draft and all authors reviewed and edited the final manuscript.

### COMPETING FINANCIAL INTERESTS

R.Y and D.S.P. are listed as inventors of the following patent application: “Devices, apparatus and method for providing photostimulation and imaging of structures” (US20110233046 A1).

## SUPPLEMENTARY INFORMATION

### Supplementary Figures S1∼S5

**Figure S1.**
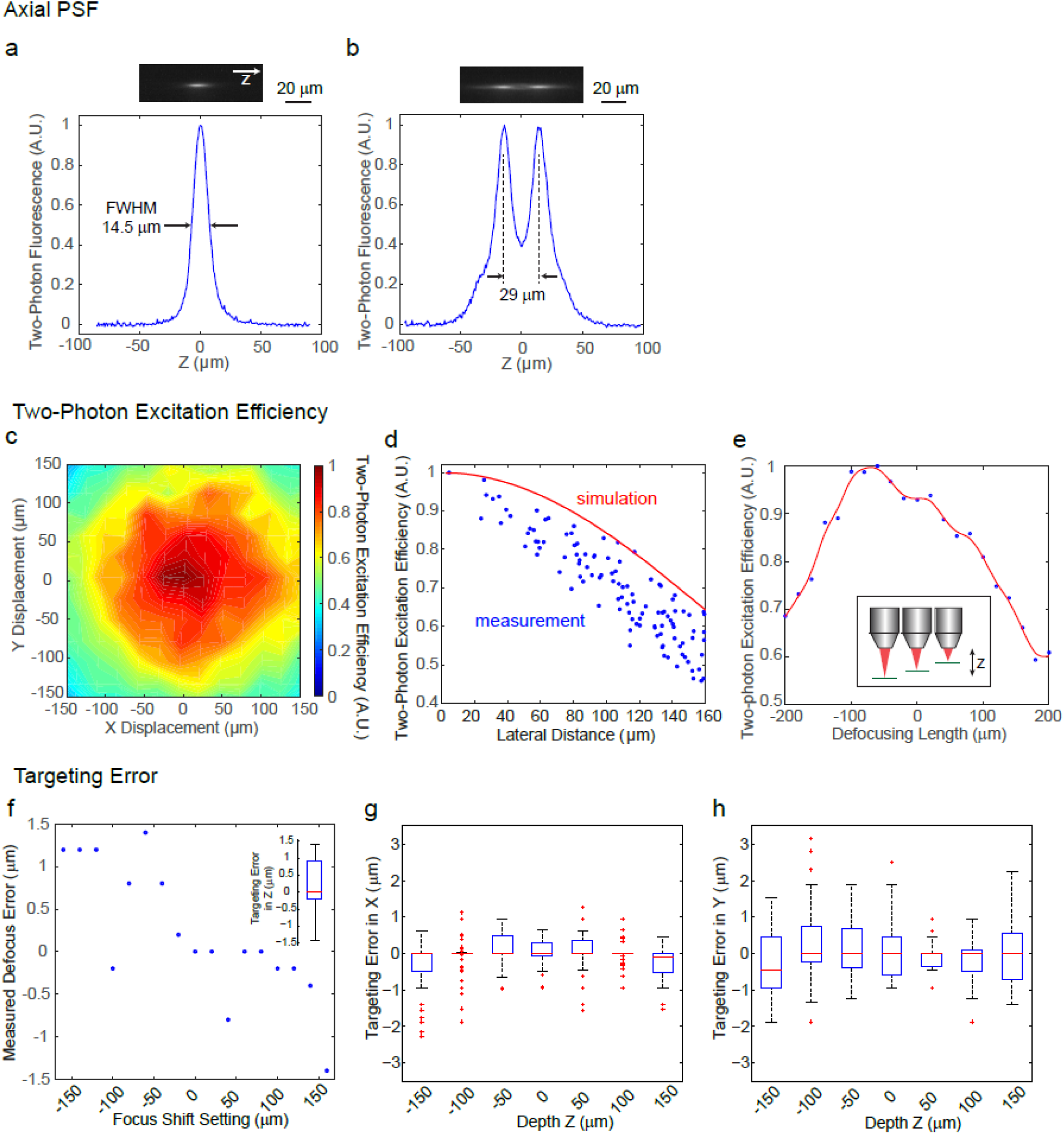
System characterization of the spatial light modulator (SLM) in the 3D microscope. (a) Measured point spread function (PSF) in the axial (*z*) direction for two-photon excitation. The FWHM is 14.5 μm, corresponding to an NA ∼ 0.35. (b) Measured axial profile of a two-photon hologram imaging where two spots was separated in 29 μm in *z*. (c) Measured SLM two-photon excitation efficiency versus lateral deflection (*x*, *y*) in the imaging plane. (d) Simulated SLM two-photon excitation efficiency versus lateral deflection in the imaging plane (red curve), with measured data (blue dot) from (c). (e) Measured SLM two-photon excitation efficiency versus defocusing length. The measured value (blue dot) is spline-fitted (red curve). (f) Measured SLM axial targeting error versus axial focus shift. Inset, boxplot of axial targeting error. Overall, the axial targeting error (absolute value) is 0.59±0.54 μm across the axial range of 300 μm. (g)-(h) Measured SLM lateral (*x*, *y*) targeting error versus axial focus shift. Overall, the lateral targeting error (absolute value) is 0.82±0.65 μm across the 3D field of view (FOV) of 240x240x300 μm^3^. In the boxplot, the central mark indicates the median, and the bottom and top edges of the box indicate the 25th and 75th percentiles, respectively. The whiskers extend to the most extreme data points (99.3% coverage if the data are normal distributed) not considered outliers, and the outliers are plotted individually using the ‘+’ symbol.

**Figure S2.**
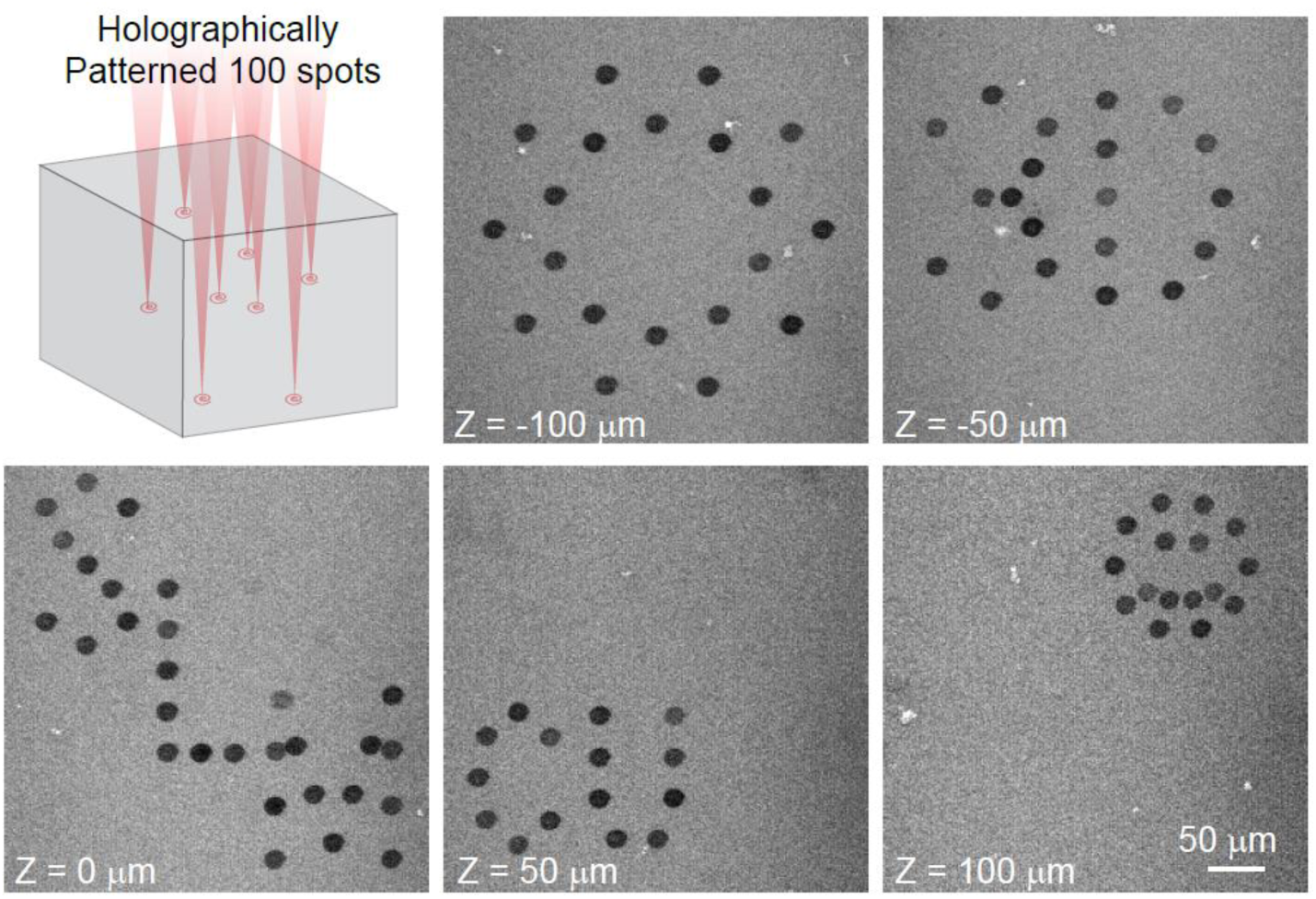
100 simultaneous spot holographic pattern spirally scanned by a post-SLM galvanometric mirror, bleaching an autofluorescence plastic slide across 5 different planes. The images show the patterns at different planes.

**Figure S3.**
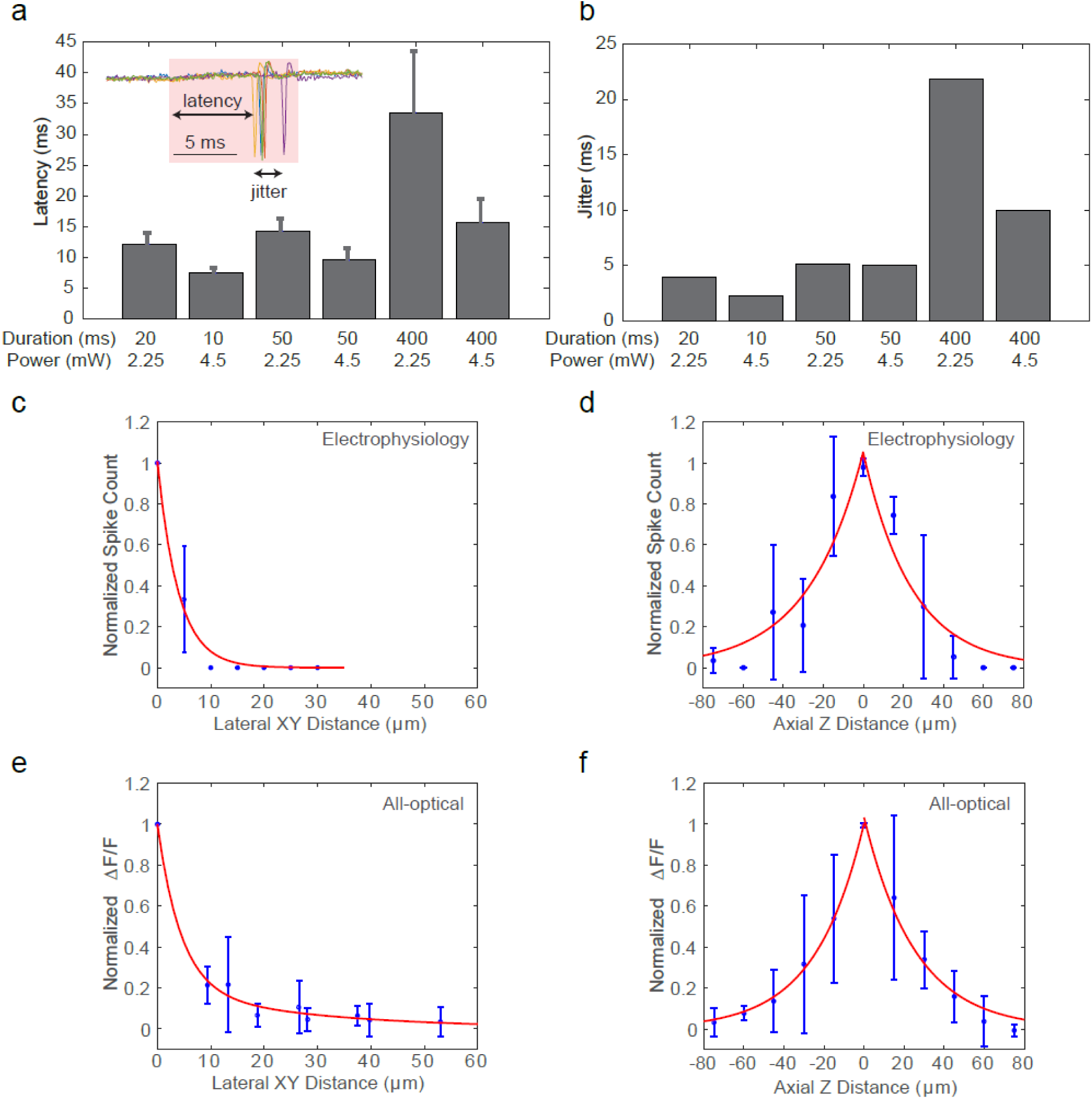
Single cell photostimulation. (a)-(b) Latency (a) and jitter (b) of a neuron evoked by photostimulation (5 trials) with different spiral duration and average laser power. The inset shows the cell-attached recording of a 10 ms spiral stimulation over 5 trials. The red bar area indicates the photostimulation period. (c)-(d) Normalized spike count versus the (c) lateral and (d) axial displacement between the centroids of the photostimulation spiral pattern and the cell body, measured by *in vivo* cell-attached electrophysiology (4 cells over 2 mice). (e)-(f) Normalized ΔF/F versus the (e) lateral and (f) axial displacement between the centroids of the photostimulation spiral pattern and the cell body, measured by *in vivo* calcium imaging [5 cells over 2 mice for (e) and 4 cells over 2 mice for (f)]. Error bars are standard error of the mean.

**Figure S4.**
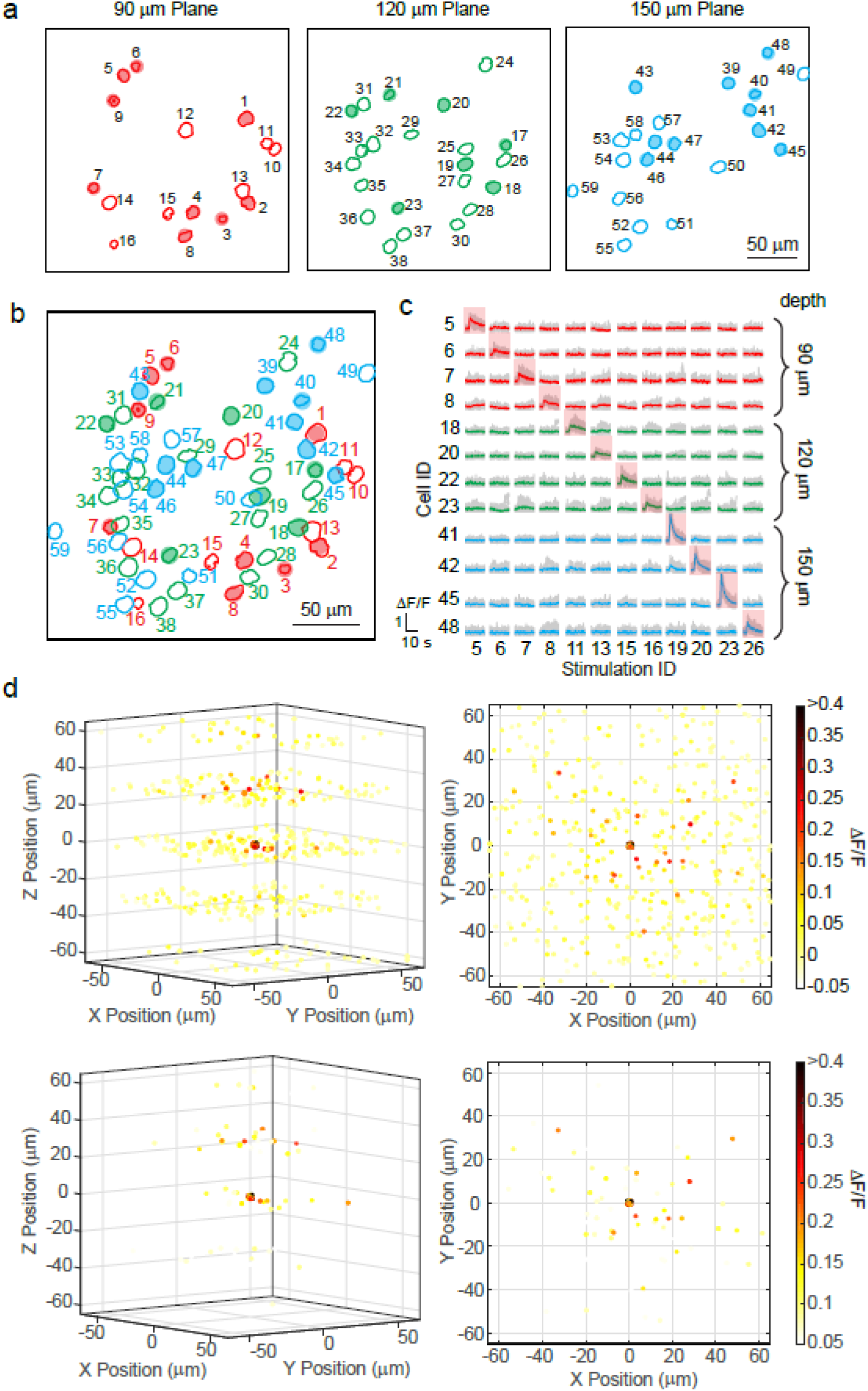
Sequential single pyramidal cell photostimulation in layer 2/3 of mouse V1 *in vivo*. (a) Contour maps showing the spatial location of the cells in three individual planes (90 μm, 120 μm, and 150 μm from pial surface). Cells with shaped color are the targeted cells. (b) 2D overlap projection of the three planes in (a). (c) Representative photostimulation triggered calcium response of the targeted cells and non-targeted cells, for different single cell photostimulation conditions. (d) Neuronal calcium response during different single cell photostimulation conditions. The spatial locations of the cells are relative to the targeted cells, which are set at the (0, 0, 0). The spatial locations of different set of conditions are randomly dithered by <1 μm in x, y, z such that the target cells do not appear to completely overlapped at (0, 0, 0). The ΔF/F response is color coded. The top and bottom panel uses two different color scales. The top panel illustrates all the cells, and the bottom panel highlights the cells showing high response. The left panel shows the 3D perspective, the right panel shows the projection in xy plane.

**Figure S5.**
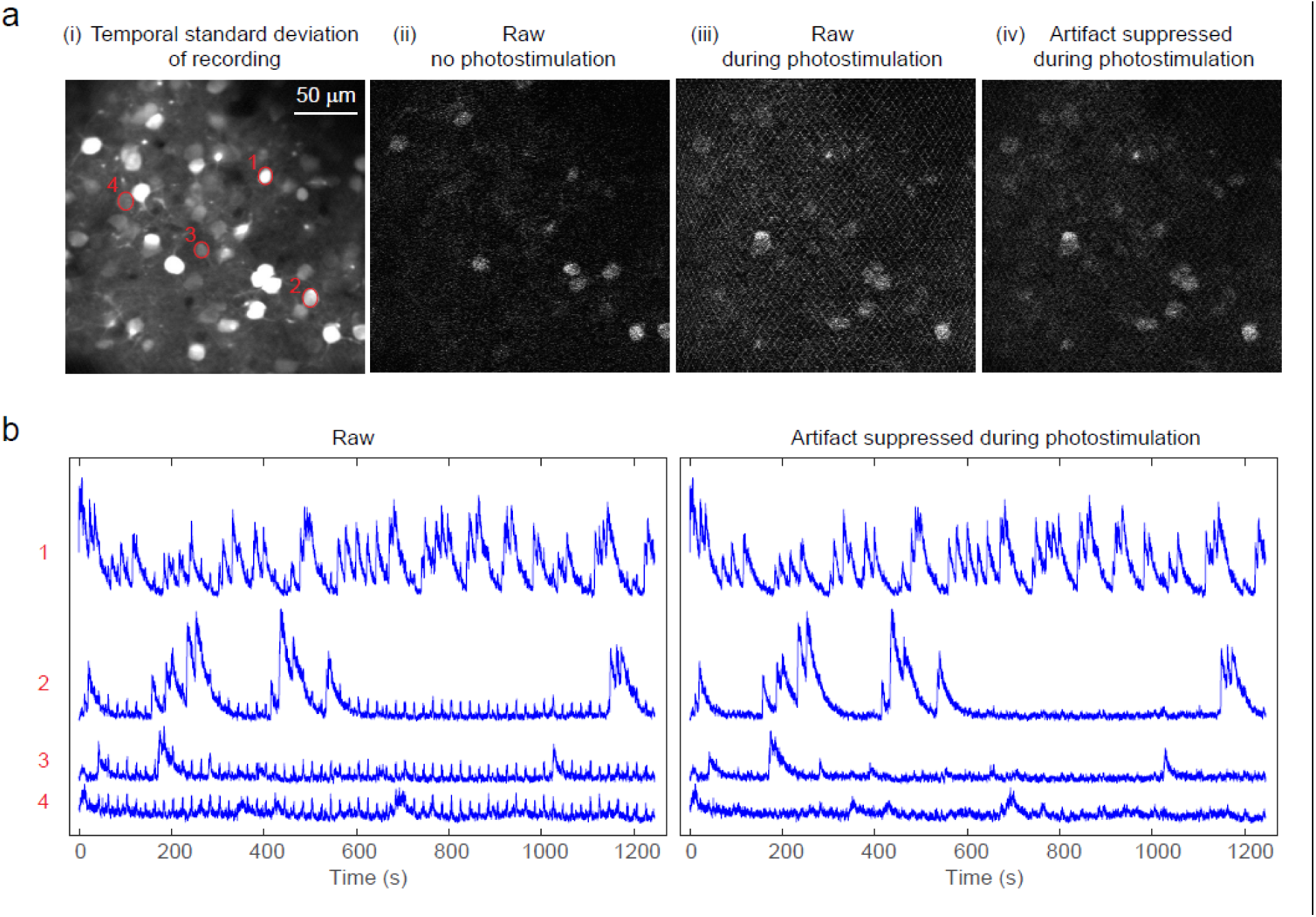
Cross talk from photostimulation laser into imaging. (a) Simultaneous calcium imaging and photostimulation in an awake mouse V1, layer 2/3. Panel i, temporal standard deviation of the recorded movie. Panel ii, a raw image frame with no photostimulation. Panel iii, a raw imaging frame during photostimulation (90 mW on sample surface). The mesh pattern comes from the stimulation artifact. Panel iv, the same image frame from panel iii but with artifact suppression by data pre-processing. (b) Representative fluorescence traces of four cells [marked in (a), with different signal-to-noise ratio] from the raw recording and that after artifact suppression.

### SUPPLEMENTARY NOTES

#### Low repetition rate pulsed-amplified laser for patterned photostimulation

In two-photon patterned photostimulation, the laser beam is split into multiple (*M*) beamlets, each of which targets an individual cell. If the total peak power of the laser is *P*_*peak*_, each cell could receive a peak power of (*P*_*peak*_/*M*). The two-photon excitation is thus scaled with (*P*_*peak*_/*M*)^2^. On the other hand, the average power *P*_*ave*_ of the laser is scaled with *P*_*peak*_\**τp*f_rep_*, where *τp* is the laser pulse width, and *f*_*rep*_ is the pulse repetition rate. A high *P*_*ave*_ could cause excessive heating on the mice brain, leading to cell damage^1^. In general, *P*_*ave*_ should not exceed 250 mW for continuous illumination.

To simultaneously photostimulate a large number of cells (a large *M*), a high *P*_*peak*_ is needed so as to maintain the required two-photon excitation for each cell. One can reduce *f*_*rep*_. The two-photon photostimulation laser we used has a low *f*_*rep*_ (200 kHz ∼ 1 MHz), about 400x ∼ 80x reduction compared to the typical *f*_*rep*_ of 80 MHz. This leads to a 400x ∼ 80x increase in *P*_*peak*_ and thus the number of possible simultaneously targeted cells *M*, with the same *P*_*ave*_ and *τp*.

The typical average power to photostimulate each cell in our experiments is 2 mW∼ 5 mW (1 MHz pulse repetition rate). We show an action potential can be evoked in a cell by ∼2.25 mW with 20 μs spiral scan duration (Fig. 1e). With ∼ 250 mW average power (below the thermal damage limit), it is estimated that ∼110 cells can be photoactivated in 20 ms, and thus ∼ 5000 cells in 1 sec (by switching between different SLM patterns; switching time < 3 ms^2^).

We note that because most opsins open ion channels, a single photoexcitation leads to multiple ions entering the cell. The average open time is also much longer than the laser’s interpulse interval (1/*f*_*rep*_). This is in contrast to fluorescence, where at most a single photon is emitted for each absorption, and the lifetime is significantly shorter than the interpulse interval. Thus opsins are ideal targets for low-repetition rate, high peak power excitation.

#### Scanning and scanless approach for photostimulation

We combine the SLM hologram and spiral scan as a hybrid strategy for patterned photostimulation on multiple cells across a 3D volume. Two alternative scanless approaches exist: pure 3D hologram and another hybrid method combining holographic patterning and temporal focusing. The former approach directly generates the full 3D hologram covering the cell bodies of targeted cells all at once. Though the simplest, the full 3D hologram has a very poor axial resolution^3^, and is thus highly subject to light contamination to the non-targeted cells. Temporal focusing^4,5^ solves this issue by coupling the holographic pattern to a grating^3^. For 3D stimulation, it could require two SLMs: one to split the laser at the lateral direction and one to adjust their focal depth^6^. Alternatively, recent report shows it is possible to use a single SLM but at a tradeoff of creating a secondary focus^7^. Regardless of the exact implementation, scanless approaches require higher laser powers per cell in general. The area-activation of scanless activation generally gives lower latencies and jitter, sometimes an important parameter in experiments, compared to scanning strategies. However, as we show, even with low powers, we have can have latencies under 10 ms, with little jitter. We expect the different photostimulation strategies complement each other, and each could be more or less advantageous depending on the exact applications. The same can be said for temporal focusing strategies. The use of temporal focusing further decouples axial from lateral extent, and helps confine the excitation PSF. This comes with added complexity, and requires more significant changes to the microscope’s optical train than our methods. Here, each discrete point in the hologram typically maintains sufficient axial confinement, even at the reduced NA used in this study. Taken together, the spiral scan strategy we adapted requires a lower laser power budget per cell, and is very scalable towards activating large number of simultaneously targeted cells, making it an advantage important tool to study ensembles in neural circuits.

#### Cross talk between imaging and photostimulation

There are two different types of cross talk in all-optical methods. The first type affects neuronal excitability, and is the result of possible photostimulation by the imaging laser. Though the C1V1 we used is a red-shifted opsin, it can still be excited at 920 ∼ 940 nm, the typical wavelength to image GCaMP6s. This cross-talk will highly depends on the expression of the calcium indicators and opsin^8,9^, and in general, the imaging laser power should be as low as possible. In the case that the calcium indicator is weakly expressed, the increased imaging power could bias the neuronal excitability. In the future, as red indicators keep improving, we may see a switch toward “blue” opsins again, as the spectral overlap between opsin and indicator can be reduced.

The second type of the cross talk affects high fidelity recording of neural activity, and is caused by fluorescence generated by the photostimulation laser, which could cause background artifact on the calcium signal recording during photostimulation. In our experiments, we use a tight bandpass filter (passband: 500 nm ∼ 520 nm) for the GCaMP6s signal detection. The C1V1 is co-expressed with mCherry, which has negligible fluorescence at the filter’s passband. On the other hand, GCaMP6s can still be excited at the photostimulation laser’s wavelength at 1040 nm. Typically this fluorescence is weak and would not impact the data analysis. However, if GCaMP6s is over-expressed and the number of simultaneously targeted neurons is high, it could cause significant contamination (background artifact) to the calcium signal recording. Since the pixel rate (∼8.2 MHz) of the calcium imaging is much faster than the photostimulation laser’s pulse repetition rate (200 kHz ∼ 1 MHz), this background artifact appears to be a mesh grid shape in the calcium imaging movie (supplementary Fig. S5). In the case that the artifact is strong, the calcium imaging movie is pre-processed so that the mesh grid shape background is replaced by their adjacent pixel value (see Methods). This pre-processing significantly suppresses the artifacts while maintaining the original signal. Nevertheless, to avoid any analysis bias, the neuronal response can be further approximated by using the ΔF/F signal just after the photostimulation, when there was no background artifact. An alternative method is to blank the PMT, or the PMTs output during the photostimulation pulse, thought this requires dedicated additional electronics. Regardless, there will be “lost” signal, and this can be treated similarly by filling in the data with interpolation.

